# Lipid Phase Behavior of the Curvature Region of Thylakoid Membranes of *Spinacia oleracea*. Isotropic Phase around the CURT1 Protein

**DOI:** 10.1101/2024.11.12.623231

**Authors:** Kinga Böde, Andrea Trotta, Ondřej Dlouhý, Uroš Javornik, Virpi Paakkarinen, Hiroaki Fujii, Ildikó Domonkos, Ottó Zsiros, Janez Plavec, Vladimír Špunda, Eva-Mari Aro, Győző Garab

## Abstract

Thylakoid membranes (TMs) of oxygenic photosynthetic organisms are flat membrane vesicles, which form highly organized, interconnected membrane networks. In vascular plants, they are differentiated into stacked and unstacked regions, the grana and stroma lamellae, respectively; they are densely packed with protein complexes performing the light reactions of photosynthesis and generating a proton motive force (pmf). The maintenance of pmf and its utilizations for ATP synthesis require sealing the TMs at their highly curved regions (CRs). These regions are devoid of chlorophyll-containing proteins but contain the curvature inducing CURVATURE THYLAKOID1 (CURT1) proteins, and are enriched in lipids. Because of the highly curved nature of this region, at the margins of grana and stroma TMs, the molecular organization of lipid molecules is likely to possess distinct features compared to those in the major TM domains. To clarify this question, we isolated CR fractions from *Spinacia oleracea* and, using BN-PAGE and western blot analysis, verified that they are enriched in CURT1 proteins and in lipids. The lipid phase behavior of these fractions was fingerprinted with ^31^P-NMR spectroscopy, which revealed that the bulk lipid molecules assume a non-bilayer, isotropic lipid phase. This finding underpins the importance of the main, non-bilayer lipid species, monogalactosyldiacylglycerol, of TMs in their self-assembly and functional activity.

## Introduction

The thylakoid membranes (TMs) exhibit a complex architecture that accommodates the main protein components performing the light reactions of oxygenic photosynthesis. In chloroplasts they form extended multilamellar networks of flattened vesicles, separating the inner and outer aqueous phases, the lumen and the stroma, respectively. In vascular plants, TMs are organized into two main domains: the granum, consisting of tightly stacked, membranes, and the stroma lamellae, which are unstacked membranes wounding around the grana in a quasi-helical manner [1, 2]. The protein composition in plant TMs displays lateral heterogeneity [3-5]. Photosystem II (PSII) and its associated light-harvesting antenna complex (LHCII) are predominantly located in the appressed membranes, while Photosystem I (PSI) and its light-harvesting antenna, LHCI, along with the ATP synthase, reside in the stroma lamellae. The cytochrome *b6f* complexes are distributed uniformly throughout the membrane system [4].

In addition to these two basic building units of TMs, a third domain, the granum margin (GM), has been proposed to be a compositionally, structurally, and functionally distinct region [6, 7]. However, different experimental techniques yielded varying interpretations of this membrane domain. On one hand certain electron microscopy (EM) studies identified the GM as highly curved areas at the periphery of grana, comprising roughly 5-7% of the TM area, while experiments employing mechanical fragmentation isolated a 60 nm wide margin annulus, revealing a significant contact area between PSII and PSI domains [6]. Recently a detailed biochemical analysis using digitonin solubilization of Arabidopsis TMs, has demonstrated that GM segregates in two distinct regions: (i) the connecting domain (CD), which physically connects grana with stroma lamellae and hosts both photosystems, and (ii) the domain containing curvature particles, which are deficient in photosystems but are enriched in CURT1 proteins [5]. This latter, highly purified CURT1-enriched domain, the curvature region (CR) fraction, has been identified using low concentration of digitonin (0.4% or 0.25%); this fraction (aka loose pellet) is largely devoid of chlorophyll-binding protein complexes [8]. Hence, CR represents a subdomain of GM. It is noteworthy that CR possesses approximately three times more lipids relative to chlorophyll (Chl) and about twice as much relative to protein than TMs [9]. This suggests that the generation of CR domain, besides CURT1, depends on the organization of the lipid molecules.

Using mainly ^31^P-NMR spectroscopy, a technique for fingerprinting the lipid phase behavior of phospholipid-containing assemblies in vivo and in vitro [10], it has been documented that the bulk lipid molecules in plant TMs display marked polymorphism: in addition to the bilayer or lamellar (L) phase, they contain an inverted hexagonal (H_II_), and at least two isotropic (I) lipid phases [11]. The non-bilayer propensity of TMs is evidently due to the presence of their main (∼50%), non-bilayer lipid species, monogalactosyldiacylglycerol (MGDG). It has also been clarified that the lipid polymorphism in isolated granum and stroma TMs is very similar to that in TMs, showing that the strikingly different protein composition of these domains does not bring about different lipid polymorphism [12]. Experiments using lipases and proteases have revealed that the non-bilayer lipid phases are found in different subdomains of TMs, distinct from the bilayer domains enriched in PSI and PSII supercomplexes. I phases were found in the lumenal compartment, where they form VDE:lipid assemblies [13], and were also proposed to be involved in membrane fusion [14]. (VDE, violaxanthin de-epoxidase, a water-soluble lipocalin-like photoprotective enzyme, the activity of which has been shown to require non-lamellar lipid phase [15].) Trypsin-digestion experiments led to the inference that lipids assuming H_II_ phase are associated with stroma-exposed proteins or polypeptides [16]. It is noteworthy, that CURT1 proteins were shown to be susceptible to trypsin digestion [7].

The aim of this work was to clarify the lipid phase behavior of CR region isolated from *Spinacia oleracea*. To this end, first we have established that our CR preparation is enriched in CURT1 protein. By using ^31^P-NMR spectroscopy we have revealed that the bulk lipid molecules in CR do not form a bilayer (L) phase but are assembled into isotropic phases.

## Materials and Methods

### Isolation of the curvature region

To isolate the curvature region (CR), or loose pellet, we followed the step-by-step protocol described by [17]. *Spinacia oleracea* (hereafter spinach) leaves were homogenized in ice-cold B1 buffer (50 mM HEPES/KOH (pH 7.5), 330 mM sorbitol, 5 mM MgCl_2_, 2.5 mM ascorbate, 0.05% (w/v) BSA), then filtered through two layers of nylon mesh and centrifuged at 5,000 × *g* at 4 °C for 4 min. Then, we suspended the resulting sediment in 30-40 mL of B2 buffer (50 mM HEPES/KOH (pH 7.5), 5 mM sorbitol, 5 mM MgCl_2_) and centrifuged again at 5,000 × *g* for 4 min. We suspended the sediment in 30-40 mL of B4 buffer (50 mM HEPES/KOH (pH 7.5), 100 mM sorbitol, 5 mM MgCl_2_), which was followed by another centrifugation with the same parameters, then we suspended again the sediment in a small amount of B4 buffer. The volume was adjusted to 0.5 mg Chl mL^-1^ with B3 buffer (15 mM Tricin (pH 7.8), 100 mM sorbitol, 10 mM NaCl, 5 mM MgCl_2_) and digitonin (Sigma-Aldrich, Burlington, MA, USA) at a final concentration of 0.4% (w/v). The suspension was stirred for 8 min with a magnetic stirrer at room temperature and then centrifuged at 1000 × *g* for 3 min. The supernatant was centrifuged for 30 mins at 10,000 × *g* to separate the core of grana; the supernatant was centrifuged at 40,000 × *g* for another 30 min and the pellet containing the CD was collected. Following these steps, the supernatant was centrifuged at 144,000 × *g* for another 1 h to separate the solid sediment (containing the stroma lamellae) and the loose pellet containing the CR. The fractions collected were suspended in B4 buffer; and the suspension was stored at -80 °C until use. *Arabidopsis thaliana* (hereafter Arabidopsis) TM and CR, obtained with 0.4% DIG, were isolated with the same procedure as in [5].

### Enzymatic treatments with wheat germ lipase and trypsin

Isolated CR particles were treated with wheat germ lipase (WGL) – a substrate nonspecific general tri-, di-, and monoglyceride hydrolase/lipase [18]. WGL was purchased from Sigma-Aldrich (Burlington, MA, USA) and was applied in 20 U mL^−1^ concentration; the treatments and incubations were performed at 5 °C. Earlier, using thin layer chromatography, we have verified that WGL digests the main TM lipid species, MGDG [16, 19].

For trypsin (T8003; trypsin from bovine pancreas; Sigma-Aldrich, Burlington, MA, USA) treatment, a stock of 300 mg mL^-1^ was dissolved in demineralized water, and 33 μL per mL of sample was added for a total concentration of 10 mg mL^-1^. The suspension was thoroughly mixed and kept at 5 °C until the start of the measurements.

### lpBN-PAGE and western blotting

Analysis of the pattern of TM protein complexes was performed by large pore Blue-Native Polyacrylamide Gel Electrophoresis (lpBN-PAGE) as described in [20]. Analysis of the amount of PSI, PSII, and CURT1 proteins was performed by immunoblotting as described in [5], by using PsaB antibody (for PSI, Agrisera www.agrisera.com catalogue numbers AS10695), CP47 antibody (kind gift of prof. Roberto Barbato) and CURT1A antibody [7]. Samples were loaded on chlorophyll basis (5 μg for lpBN-PAGE and 1 μg for immunoblotting).

### Scanning electron microscopy

The membrane fractions were fixed in 2.5% glutaraldehyde for 2 hours, settled on poly-L-lysine – coated polycarbonate filter for 45 min. After post-fixation in 1% OsO_4_ for 50 min, the samples were dehydrated in aqueous solutions of increasing ethanol concentrations, critical point dried, covered with 5 nm gold by a Quorum Q150T ES (Quorum Technologies, Lewes, UK) sputter and observed in a JEOL JSM-7100F/LV (JEOL SAS, Croissy-sur-Seine, France) scanning electron microscope.

### ^31^P–NMR spectroscopy

^31^P-NMR spectroscopy tracks the motion of bulk phosphatidylglycerol (PG) molecules in the TM, serving as a sensitive internal probe of the lipid phase behavior. This is justified by the fact that PG displays no lateral heterogeneity among the bulk phase of TM domains [21] and is homogenously distributed in TM lipid mixtures [22].

^31^P-NMR spectroscopy was performed as described in [12]. Briefly, the spectra were recorded at 5 °C, using an Avance Neo 600 MHz NMR spectrometer (Bruker, Billerica, MA, USA) with a BBFO SmartProbe that was tuned to the P-atom frequency. The sample was loaded into 5 mm diameter NMR tubes. For spectra acquisition, 40° RF pulses with an inter-pulse time of 0.5 s were applied without ^1^H-decoupling, as in earlier experiments [23]. As an external chemical-shift reference 85% solution of H_3_PO_4_ in water was used. In the saturation transfer (ST) experiments, we employed RF pulses with low power at the designated frequency for a duration of 0.3 s, followed by 40° pulses with an acquisition time of 0.2 s and a repetition time of 0.5 s. The intensity of the pre-saturation pulse was adjusted based on the intensity of the saturated peak. For the pre-saturation RF pulses, the field strength was set to 40 Hz.

CR samples from two batches, with very similar features, were averaged to improve the signal-to-noise ratio. For ^31^P-NMR data processing, TopSpin software (Bruker, Billerica, MA, USA) was used. The figures were plotted using MATLAB R2020b (MathWorks, Inc., Portola Valley, CA, USA).

### Fourier-transform infrared spectroscopy

To determine the lipid-to-protein ratios in TMs and CR, we used FTIR spectroscopy (see Supplementary Material). Membrane fraction suspensions were washed in D_2_O-based PBS (phosphate-buffered saline) solution for complete H_2_O to D_2_O exchange. The sample was layered between CaF_2_ windows, separated by an aluminum spacer, and placed in a Bruker Vertex70 FTIR spectrometer. Spectra were recorded at 5 °C between 4,000 and 900 cm^-1^, 512 interferograms were collected for each spectrum, the spectral resolution was 2 cm^-1^. The infrared absorption spectrum of the samples was calculated from the background and sample of single beam spectra with Opus software of Bruker. For the analysis of the structural properties of the membrane, the ester C=O plus the amide I region, which in this paper will be referred to as ‘Ester + Amide I’ region was used between 1,800 and 1,595 cm^-1^. To obtain the relative intensities of the ‘ester’ and the ‘amide I’ bands in the ‘Ester + Amide I’ region, a 3rd order polynomial was fitted and subtracted as baseline. The area under the curves were calculated by using in-built MATLAB functions, after trimming the spectral regions corresponding to the ‘ester’ and ‘amide I’ bands.

## Results

### Pattern of protein complexes in TM fractions obtained by digitonin solubilization

The pattern of the protein complexes in spinach TM and the fractions obtained by 0.4% digitonin solubilization were analyzed by lpBN-PAGE (Figure 1A). The typical pattern observed in fractions obtained with the same procedure in Arabidopsis was observed [5]. In particular, the 10k fraction, enriched in grana core, contained the majority of the PSII-LHCII supercomplexes (sc) and LHCII complexes (M-, L- and monomeric LHCII). On the other hand, the 144k fraction showed the typical pattern of stroma lamellae, being enriched in PSI-LHCI and other PSI complexes, and depleted in LHCII. The 40k fraction, enriched in CD, had somewhat intermediate features. Considering the three strong pellet fractions (i.e. 10k, 40k and 144k), PSII monomer and Cyt b_6_f were mostly enriched in 10k and 40k fractions. However, the highest amount of PSII monomer and Cyt b_6_f, together with ATP synthase, was present in the CR fraction, which was instead strongly depleted of Chl-binding protein complexes (Figure 1A). A similar pattern was visible in the Arabidopsis CR which, in line with expectations using 0.4% digitonin, contained Chl-binding protein complexes to some extent (compare with [5]).

**Figure 1.**
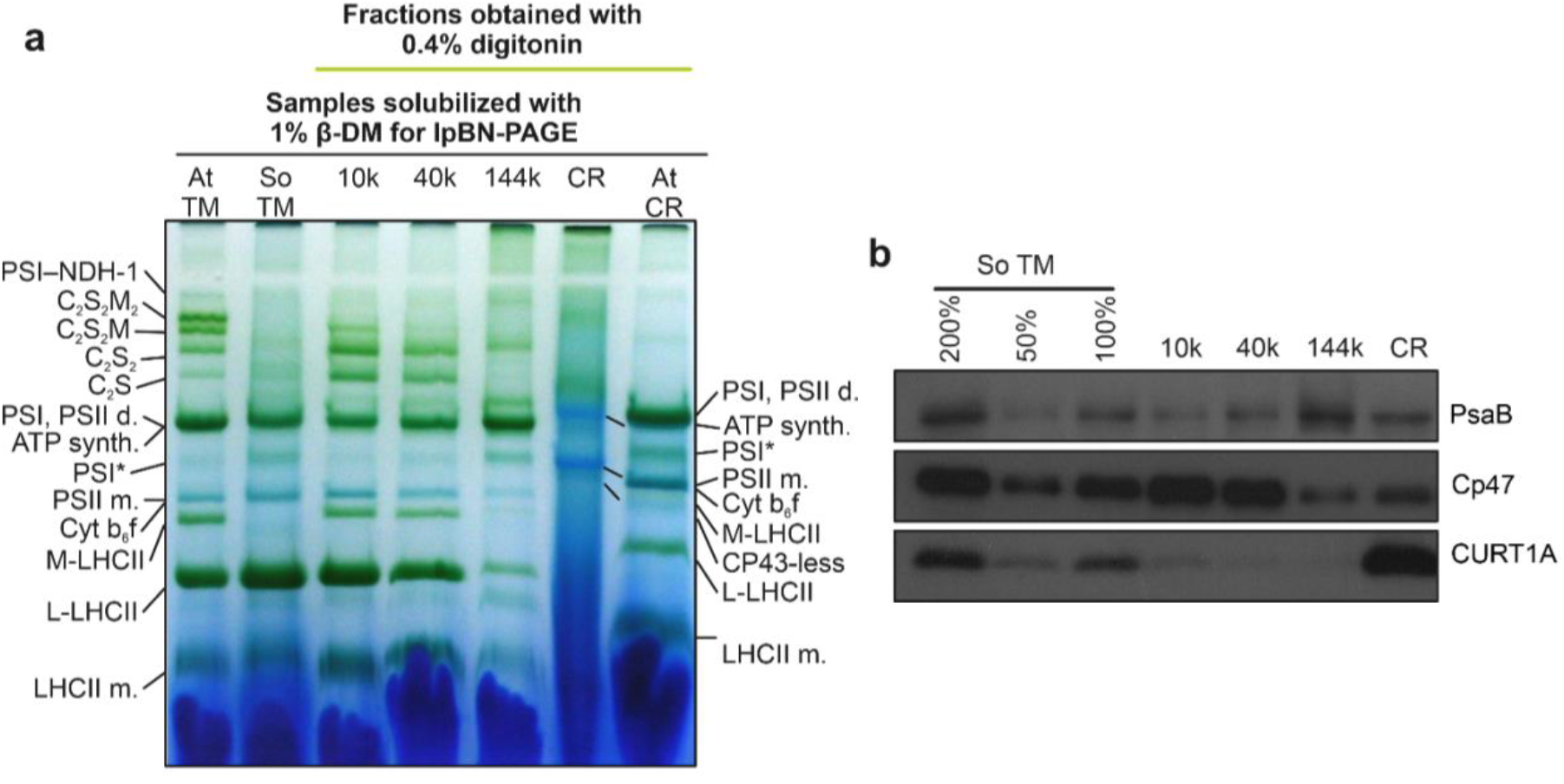
(**A**), lpBN-PAGE analysis of TM and fractions after solubilization with 0.4% digitonin of spinach (So) TM. At: Arabidopsis thaliana. (**B**) immunoblot with PsaB, Cp47and CURT1A antibodies of the same TM and fractions as in A. A cross-reactivity scale with So TM is shown on the left.

The analysis by immunoblotting confirmed the results obtained with lpBN-PAGE (i.e. enrichment of PSII in 10k and of PSI in 144k (Figure 1B). Consistent with previous results [5, 8] the CR fraction had the highest enrichment in CURT1, indicating that, besides other proteins collected in the loose pellet due to digitonin solubilization, this was the fraction where most of the curvature domain of the GM was collected.

### Morphology of curvature region

For the structural characterization of CR, scanning electron microscopy (SEM) was performed (Figure 2). CR fraction consisted of small particles with a tendency of creating large self-aggregates and/or smaller vesicle-like assemblies.

**Figure 2.**
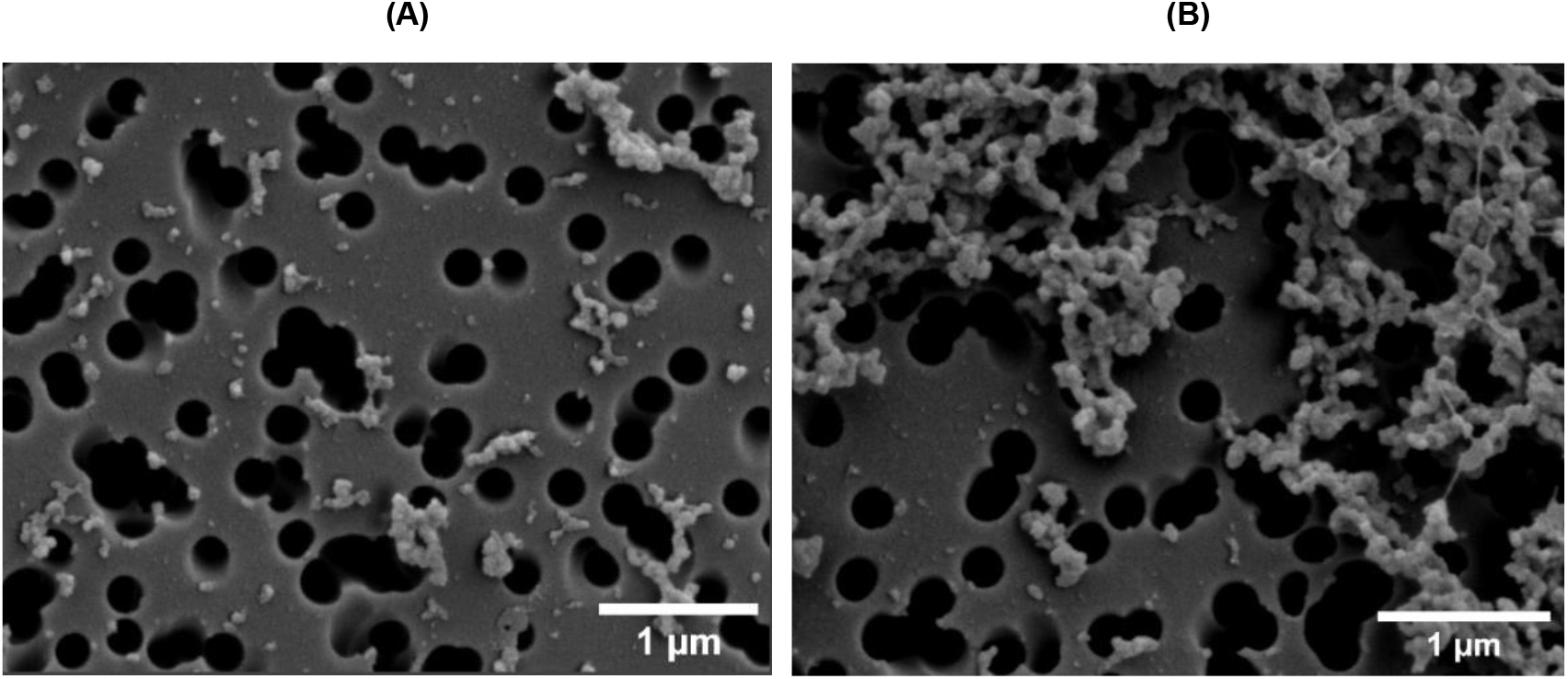
SEM micrographs of CR fractions showing bent membrane particles (**A**), which often coagulate (**B**).

These structures are distinct from those observed in the grana and stroma fractions, where membrane vesicles appear predominantly, as reported in previous studies [9, 24]. Notably, the morphology of these particles aligns well with the observations of [9] using TEM.

### Lipid phase behavior of CR particles

As evidenced by the ^31^P-NMR spectra shown in Figure 3, the CR particles do not contain L and H_II_ phases, which are present in intact TMs and the granum and stroma subchloroplast membrane particles isolated with the same procedure [12] (see also the ^31^P-NMR spectrum of stroma lamellae in the Supplementary Material). Instead, the spectra of CR preparations are dominated by well-discernible I phases with peak positions at around 0.15 and 2.7 ppm. To probe the spectral shapes and resolve overlapping phases, we applied low power radiofrequency pulses at selected chemical shifts [12, 23]. These saturation transfer experiments confirmed the presence of these two strongly overlapping phases, although other resonances with smaller amplitudes could also be found in the band structure.

**Figure 3.**
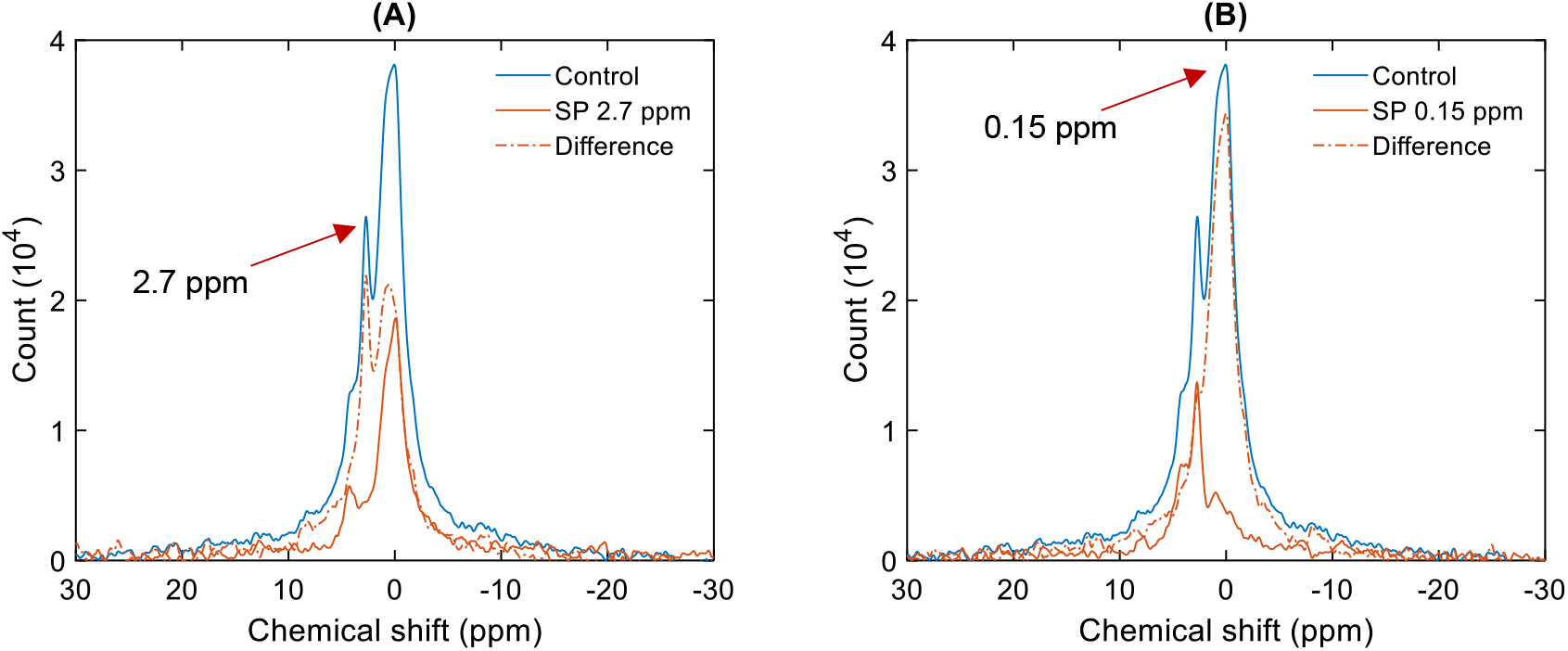
^31^P-NMR spectra of CR under control conditions, displaying also the effects of saturation pulses (SP) at 2.7 ppm (**A**) and 0.15 ppm (**B**). The number of scans was 12,800.

The resonance bands that originate from the lipid phases were sensitive to WGL digestion (Figure 4A) similarly to what has been observed in TMs and the granum and stroma subchloroplast particles [12, 16]. This shows that the character of the I phases in CR closely resembles those in intact TM and in the digitonin-fragmented grana and stroma lamellae.

**Figure 4.**
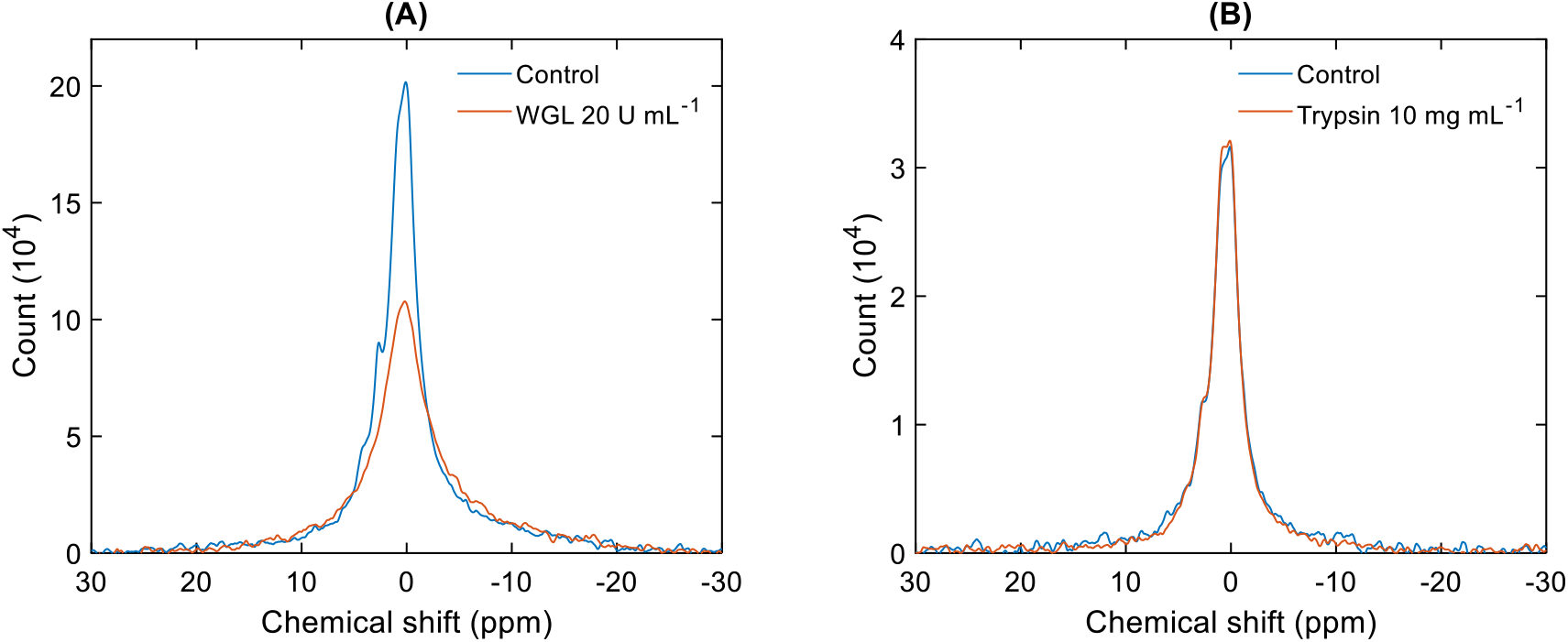
^31^P-NMR spectra of untreated (Control) and WGL-treated (20 U mL^−1^, 25,600 scans) (**A**) and trypsin-treated (10 mg mL^−1^, 19,200 scans) (**B**) CR samples.

In contrast, trypsin treatment did not elicit any sizeable effect on the spectra (Figure 4B). As we have previously shown [16], trypsin treatment selectively degraded the ^31^P-NMR detectable H_II_ phase of TMs, which revealed that this lipid phase is associated with proteins or extrinsic protein domains protruding in the stromal aqueous side of TMs. Although CURT1 has been shown to be susceptible to trypsin digestion, the absence of H_II_ phase and the trypsin-insensitive nature of the detected non-bilayer lipid phases shows that these proteins do not contribute to the formation of the H_II_ phase.

It is to be noted here, that while plastoglobuli might contribute to the composition of CR fraction [5], they display no sizeable ^31^P-NMR signal [16]; hence, they are highly unlikely to participate in the isotropic organization of the lipid molecules around CURT1 proteins.

## Discussion

In this paper, we confirmed the presence of CURT1 in the CR fraction of *Spinacia oleracea* which corroborates the notion that this protein plays the main role in generating membrane curvature in various species [7, 20, 25]. Notably, the CURT1 protein was detected using an antibody originally developed for *Arabidopsis thaliana*, demonstrating its cross-reactivity and reinforcing the evolutionary conserved nature of CURT1 across taxa.

^31^P-NMR spectroscopy – which tracks the motion of bulk PG molecules in different lipid phase environments – has unveiled that the CR fraction of TMs displays merely an intense and a weaker isotropic (I) lipid phase. The lack of the lamellar (L) phase in CR preparations is evidently the consequence of the deficiency of PSII and PSI supercomplexes, which in TMs stabilize the bilayer; similarly to that shown for LHCII:MGDG macroassemblies [26]. Note that the absence of L phase is unlikely to be caused by the usage of digitonin since both the granum and the stroma TM subdomains – with the same isolation procedure – preserve the bilayer (L) phase [12].

The presence of I phases in the CR might be attributed to curved bilayers [27]. However, the observed inverse relationship between the insulation efficiency of TMs and the intensity of I phase(s) renders this explanation unlikely; with increasing I phase contributions at the expense of L phase, the permeability of membrane increases [28]. Hence, evidently, the organization of lipid molecules in CR deviates from a typical “well-sealed” bilayer arrangement. It is interesting to note here that, as argued by [29], the lipid polymorphism of TMs is not necessarily in conflict with the efficient energization of TMs and utilization of pmf for ATP synthesis. Also, as pointed out by [30], “a membrane composed solely of lamellar lipids would be an optimum insulator, only non-compatible with cell function, i.e., life.”.

Overall, these findings strongly indicate that the lipids surrounding CURT1 proteins are organized into isotropic arrays. This means that the lipids surrounding the CURT1 protein lacks the inherent tendency to adopt a bilayer arrangement in the highly curved CR of TMs. These data underline the role of the major non-bilayer lipid, MGDG, of plant TMs in their self-assembly; and in general, corroborate the fundamental importance of lipid polymorphism in these major, energy-converting membranes.

## Acknowledgments

We are grateful to the CERIC-ERIC facilities at the Slovenian NMR Center for providing access to the ^31^P-NMR spectroscopy, as well as for the financial assistance for travel and lodging. This research was supported by the Czech Science Foundation (GAČR 23– 07744 S to G.G.), the Erkko Foundation (to E-M.A) and the European Union under the LERCO project (CZ.10.03.01/00/22_003/0000003) via the Operational Programme Just Transition. This work was supported by the Slovenian Research and Innovation Agency (ARIS, grant no. P1-0242).

## Author contributions

The study was conceptualized by G.G., K.B., A.T., E-M.A. and V.Š.; TM and CR preparations were isolated by O.Zs. and K.B.; ^31^P NMR spectroscopy measurements were performed by U.J., supervised by J.P.; data analyses were performed by K.B. and O.D., with the help of U.J., supervised by G.G.; spectral characterizations of the membranes were performed by K.B., O.D., supervised by V.Š.; FTIR measurements were carried out and analyzed by K.B.; SEM experiments were performed by I.D.; lpBN-PAGE and immunoblotting was performed by V.P. and H.F., supervised by A.T. and E-M.A. The paper was written by G.G., K.B. and A.T., with all authors contributing to the writing.

## References

1. Mustárdy, L. and G. Garab, Granum revisited. A three-dimensional model - where things fall into place. Trends in Plant Science, 2003. 8(3): p. 117–122.

2. Bussi, Y., et al., Fundamental helical geometry consolidates the plant photosynthetic membrane. Proc Natl Acad Sci U S A, 2019. 116(44): p. 22366–22375.

3. Andersson, B. and J.M. Anderson, Lateral heterogenity in the distribution of chlorophyll-protein complexes of the thylakoid membranes of spinach-chloroplast. Biochimica Et Biophysica Acta (BBA), 1980. 593(2): p. 427–440.

4. Dekker, J.P. and E.J. Boekema, Supramolecular organization of thylakoid membrane proteins in green plants. Biochim Biophys Acta, 2005. 1706(1-2): p. 12–39.

5. Trotta, A., et al., Defining the heterogeneous composition of Arabidopsis thylakoid membrane. The Plant Journal, 2025. 121(3): p. e17259.

6. Albertsson, P., A quantitative model of the domain structure of the photosynthetic membrane. Trends Plant Sci, 2001. 6(8): p. 349–58.

7. Armbruster, U., et al., Arabidopsis CURVATURE THYLAKOID1 Proteins Modify Thylakoid Architecture by Inducing Membrane Curvature. Plant Cell, 2013. 25(7): p. 2661–2678.

8. Trotta, A., et al., The role of phosphorylation dynamics of CURVATURE THYLAKOID 1B in plant thylakoid membranes. Plant Physiology, 2019. 181(4): p. 1615–1631.

9. Koochak, H., et al., The structural and functional domains of plant thylakoid membranes. The Plant Journal, 2019. 97(3): p. 412–429.

10. Watts, A., NMR of Lipids, in Encyclopedia of Biophysics, G.C.K. Roberts, Editor. 2013, Springer Berlin Heidelberg: Berlin, Heidelberg. . 1727–1738.

11. Garab, G., et al., Structural and functional roles of non-bilayer lipid phases of chloroplast thylakoid membranes and mitochondrial inner membranes. Progress in Lipid Research, 2022. 86: p. 101163.

12. Dlouhý, O., et al., Lipid Polymorphism of the Subchloroplast—Granum and Stroma Thylakoid Membrane—Particles. I. 31P-NMR Spectroscopy. Cells, 2021. 10(9): p. 2354.

13. Dlouhý, O., et al., Modulation of non-bilayer lipid phases and the structure and functions of thylakoid membranes: effects on the water-soluble enzyme violaxanthin de-epoxidase. Scientific Reports, 2020. 10(1).

14. Böde, K., et al., Role of isotropic lipid phase in the fusion of photosystem II membranes. Photosynthesis Research, 2024. 161(1-2): p. 127–140.

15. Goss, R. and D. Latowski, Lipid Dependence of Xanthophyll Cycling in Higher Plants and Algae. Front Plant Sci, 2020. 11: p. 455.

16. Dlouhý, O., et al., Structural Entities Associated with Different Lipid Phases of Plant Thylakoid Membranes - Selective Susceptibilities to Different Lipases and Proteases. Cells, 2022. 11(17): p. 2681.

17. Suorsa, M., et al., Light acclimation involves dynamic re‐organization of the pigment– protein megacomplexes in non‐appressed thylakoid domains. The Plant Journal, 2015. 84(2): p. 360–373.

18. Kublicki, M., et al., Wheat germ lipase: isolation, purification and applications. Critical Reviews in Biotechnology, 2021.

19. Böde, K., et al., Role of isotropic lipid phase in the fusion of photosystem II membranes. Photosynth Res, 2024. 161(1-2): p. 127–140.

20. Sandoval-Ibáñez, O., et al., Curvature thylakoid 1 proteins modulate prolamellar body morphology and promote organized thylakoid biogenesis in Arabidopsis thaliana. Proceedings of the National Academy of Sciences, 2021. 118(42): p. e2113934118.

21. Duchene, S. and P.A. Siegenthaler, Do glycerolipids display lateral heterogeneity in the thylakoid membrane? Lipids, 2000. 35(7): p. 739–744.

22. van Eerden, F.J., et al., Characterization of thylakoid lipid membranes from cyanobacteria and higher plants by molecular dynamics simulations. Biochim Biophys Acta, 2015. 1848(6): p. 1319–30.

23. Krumova, S.B., et al., Phase behavior of phosphatidylglycerol in spinach thylakoid membranes as revealed by 31P-NMR. Biochim Biophys Acta, 2008. 1778(4): p. 997–1003.

24. Dlouhý, O., et al., Lipid Polymorphism of the Subchloroplast—Granum and Stroma Thylakoid Membrane–Particles. II. Structure and Functions. Cells, 2021. 10(9): p. 2363.

25. Heinz, S., et al., Thylakoid Membrane Architecture in Synechocystis Depends on CurT, a Homolog of the Granal CURVATURE THYLAKOID1 Proteins. Plant Cell, 2016. 28(9): p. 2238–2260.

26. Simidjiev, I., et al., Self-assembly of large, ordered lamellae from non-bilayer lipids and integral membrane proteins in vitro. Proc Natl Acad Sci U S A, 2000. 97(4): p. 1473–6.

27. Cullis, P.R. and B. de Kruijff, Lipid polymorphism and the functional roles of lipids in biological-membranes. Biochimica Et Biophysica Acta (BBA), 1979. 559(4): p. 399–420.

28. Ughy, B., et al., Lipid-polymorphism of plant thylakoid membranes. Enhanced non-bilayer lipid phases associated with increased membrane permeability. Physiologia Plantarum, 2019. 166(1): p. 278–287.

29. Garab, G., et al., Lipid polymorphism of plant thylakoid membranes. The dynamic exchange model - facts and hypotheses. Physiologia Plantarum, 2025.

30. Goni, F.M., The basic structure and dynamics of cell membranes: an update of the Singer-Nicolson model. Biochim Biophys Acta, 2014. 1838(6): p. 1467–76.

