## Supplementary information for "Lipid Phase Behavior of the Curvature Region of Thylakoid Membranes of *Spinacia oleracea*. Isotropic Phase around the CURT1 Protein"

The highly curved CR has been shown to have lower protein/lipid ratio compared to the granum and stroma domains (Koochak et al. 2018), which we also confirmed in these experiments (SFigure 1). In SFigure 1, by comparing the area under the curve in the FTIR spectrum between the resonances originating from ester bonds characteristic to lipids between 1750 and 1700 cm$^{-1}$ and the amide bonds characteristic of proteins between 1700 and 1600 cm$^{-1}$, it is evident that the relative amount of lipids in the CR is higher compared to TM.

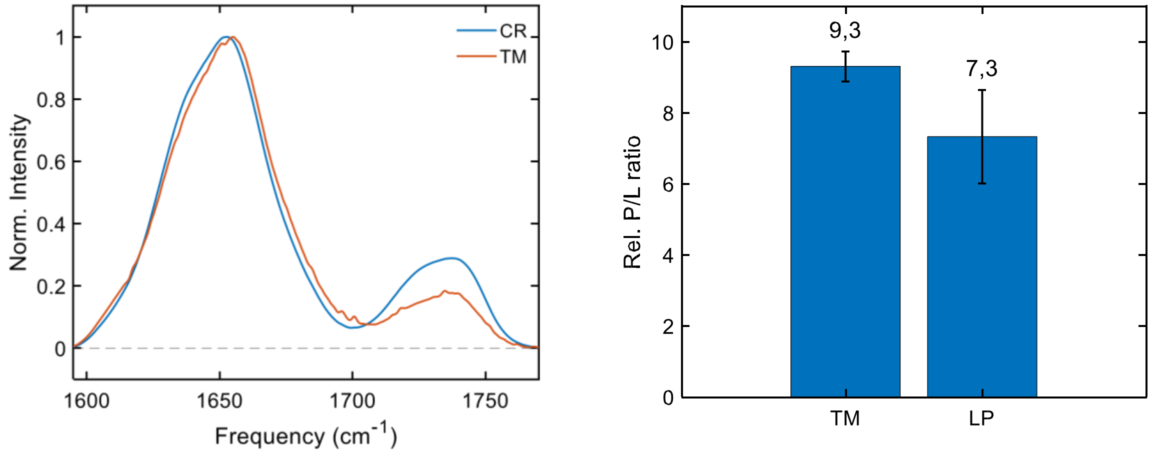


*SFigure 1: FTIR spectra of isolated TM and CR in the 'amide I' and 'ester' regions (Left), and a bar chart (Right) showing protein/lipid (P/L) ratios calculated from the integrated areas of the 'amide I' (1700-1600 cm$^{-1}$) and 'ester' (1750-1700 cm$^{-1}$) FTIR bands based on three biological replicates ± SD.*

CR particles lack the L and $H_{II}$ phases that are characteristic of intact TMs, granum and stroma subchloroplast membrane particles (Dlouhý et al. 2021a). It is important to note that while TM isolation does not require digitonin fragmentation, the granum and stroma subchloroplast particle isolation procedures use relatively high digitonin concentrations – considerably higher than applied here (see Methods). Nevertheless, the granum and stroma particles display L and $H_{II}$ phases, as their dominant lipid phases. To further substantiate that the absence of L (and $H_{II}$) phase(s) cannot be ascribed to the use of digitonin, we show that the stroma particles (144,000 x g solid sediment) from the same batch, exhibits the characteristic L and $H_{II}$ phases (Supplementary Figure 2). In particular, the averaged $^{31}$P-NMR spectrum and its mathematical deconvolution shows comparable polymorphism to data on stroma TMs published earlier (Dlouhý et al. 2021a). Thus, the lipid phases behavior appears to reflect the intrinsic feature of this CR fraction.

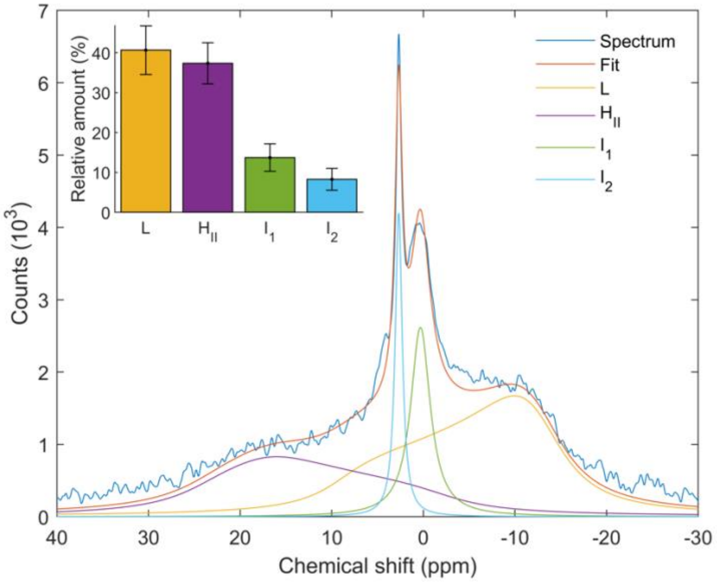


*SFigure 2: $^{31}$P-NMR spectrum of stroma thylakoids isolated from Spinacia oleracea, averaged from 3 biological replicates± SD. Number of scans 25 600. The relative amount of the lipid phases are as follows; L phase: 40.7±6.1, $H_{II}$: 37.4±5.1; $I_1$: 13.7±3.4; $I_2$:8.3±2.*

Koochak, H., Puthiyaveetil, S., Mullendore, D. L., Li, M., & Kirchhoff, H. (2019). The structural and functional domains of plant thylakoid membranes. *The Plant journal : for cell and molecular biology*, *97*(3), 412–429. https://doi.org/10.1111/tpj.14127

Dlouhý, O., Javornik, U., Zsiros, O., Šket, P., Karlický, V., Špunda, V., Plavec, J., & Garab, G. (2021). Lipid Polymorphism of the Subchloroplast-Granum and Stroma Thylakoid Membrane-Particles. I. 31P-NMR Spectroscopy. *Cells, 10*(9), 2354. https://doi.org/10.3390/cells10092354
